# Multiple independent origins of apicomplexan-like parasites

**DOI:** 10.1101/636183

**Authors:** Varsha Mathur, Martin Kolisko, Elisabeth Hehenberger, Nicholas AT Irwin, Brian S. Leander, Árni Kristmundsson, Mark A Freeman, Patrick J Keeling

**Author notes:** Corresponding authors (V.M); (M.K). VM and MK contributed equally to this work.

## Abstract

Apicomplexans are a diverse group of globally important parasites, that include pathogens like *Plasmodium*, the causative agent of malaria. Despite their current obligate parasitic nature, apicomplexans evolved from photosynthetic algae and retain a remnant plastid (chloroplast). Such a complex evolutionary transition was unexpected, but here we show that it occurred at least three times independently. Using single-cell genomics and transcriptomics from diverse uncultivated parasites, we find that two genera previously classified within the Apicomplexa, *Piridium* and *Platyproteum*, form separately branching lineages in phylogenomic analyses. Both retain cryptic plastids with genomic and metabolic features convergent with apicomplexans. These findings suggest a predilection in this lineage for both the loss of photosynthesis and the transition to a morphologically similar parasitic lifestyle, resulting in multiple lineages of highly convergent animal parasites.

## Main Text

The apicomplexans are a group of obligate animal pathogens that include *Plasmodium* (malaria), *Toxoplasma* (toxoplasmosis) and *Cryptosporidium* (cryptosporidiosis)(*1*). They are united by a complex and distinctive suite of cytoskeletal and secretory structures related to infection called the apical complex, which are used to recognize and gain entry into animal host cells. Apicomplexans are known to have evolved from free-living photosynthetic ancestors and retain a relict plastid (the apicoplast), which is non-photosynthetic, but houses a number of other essential metabolic pathways(*2*). The closest relatives of apicomplexans includes a mix of both photosynthetic algae (chromerids) as well as non-photosynthetic microbial predators (colpodellids)(*3*), and genomic analyses of these free-living relatives have revealed a great deal about how such a dramatic transition may have taken place, as well as how parasitism originates more generally(*4*).

To gain a deeper understanding of the origin of parasitism in apicomplexans, we used single-cell sequencing to characterize the genomes and transcriptomes from a number of uncultivated parasites representing poorly-studied lineages of apicomplexans. Specifically, we generated transcriptome data from individual trophozoite cells of the gregarine apicomplexans *Monocystis agilis, Lecudina tuzetae, Pterospora schizosoma, Heliospora capraellae*, and *Platyproteum* sp., using single cells documented microscopically and manually isolated directly from their animal hosts (Fig. 1A-E). In addition, we generated both genomic and transcriptomic data from gamogonic stages of *Piridium sociabile*, an apicomplexan isolated from the foot tissue cells of the common whelk, *Buccinum undatum*, which inhabits marine waters (Fig. 1F). These gregarines represent subgroups of both marine (*Pterospora, Heliospora, Lecudina* and *Platyproteum*) and terrestrial (*Monocystis*) parasites, and the limited available molecular data (from small subunit (SSU) rRNA) are divergent but generally show them to be diverse, early-branching apicomplexans (*5*–*8*). *Platyproteum* was the most recently described by detailed microscopy and molecular phylogenetic analyses using SSU rDNA sequences; these data suggest that it is a particularly deep-branching apicomplexan(*9,10*). *Piridium sociabile* is even more poorly-studied; found in 1932 as an intracellular infection and was morphologically classified as a schizogregarine (*11*).

**Fig. 1:**
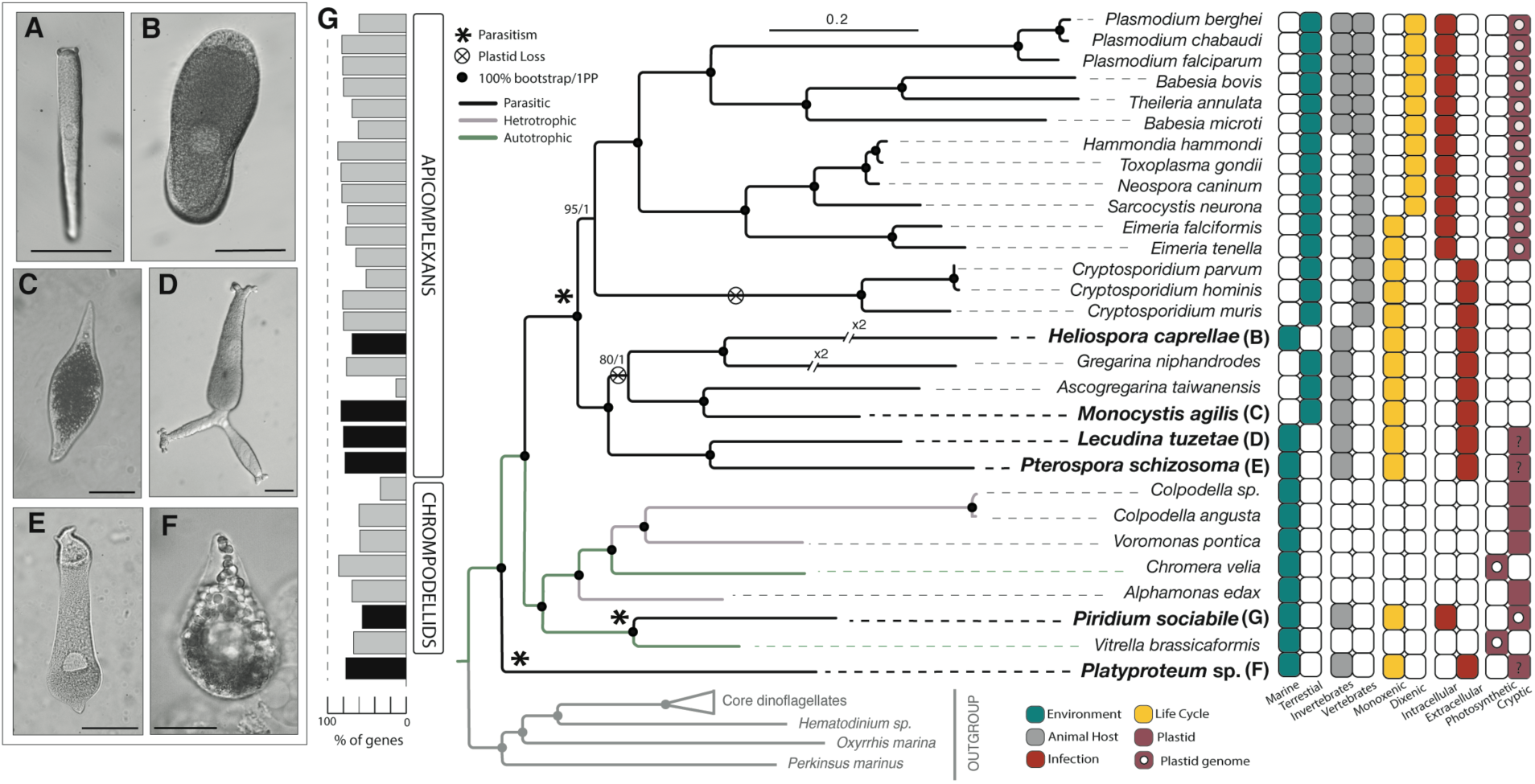
Phylogenomic tree of the Apicomplexa and relatives. Light micrographs of single-cell trophozoites are of *H. caprellae* (B) *L. tuzatae* (C) *M. agilis* (D) *P. schizosoma* and (E) *Platyproteum* sp. (scale bar = 50μm) (F) Light micrograph of a single-cell gamont of *P.sociabile* (scale bar = 15μm). (G) Maximum likelihood tree generated from an alignment comprising 198 genes and 58,116 sites under the C40+LG+Г4+F substitution model with both non-parametric bootstraps (*n* = 500) and posterior probabilities (PP) shown. Black circles represent 100% bootstrap support and 1.0 Bayesian PP, and all other support values are indicated beside the node. New transcriptomes are shown in bold lettering. The percentage of genes present in the phylogeny for each taxon are shown on the left and are shaded in black for newly sequenced transcriptomes. On the right are characters corresponding to each taxa.

The relationships of these six taxa to the Apicomplexa were examined by phylogenomics using a concatenated alignment of 39 taxa and 189 nucleus-encoded proteins that have been previously used in in both eukaryote-wide and phylum-level phylogenies (*12, 13*). Their positions in the resulting tree are strongly and consistently resolved by both maximum likelihood (C40+LG+Г4+F model) and Bayesian (CAT-GTR) analyses (Fig. 1G). Surprisingly, the phylogeny shows that neither *Piridium* or *Platyproteum* branch within the Apicomplexa. Instead, *Piridium* branches within the sister group to the Apicomplexa, the ‘chrompodellids’ (chromerids + colpodellids), with complete support as sister to the photosynthetic alga *Vitrella brassicaformis*. *Platyproteum* forms a new lineage, also with complete support, sister to the clade consisting of apicomplexans and chrompodellids collectively. The four more canonical gregarines (*Monocystis, Lecudina, Pterospora* and *Heliospora*) formed a monophyletic group of deep-branching apicomplexans that interestingly excludes *Cryptosporidium*. This robust phylogeny not only confirms that photosynthesis was lost multiple times independently around the origin of the Apicomplexa, but more surprisingly shows that the highly-derived mode of animal parasitism that is characteristic of the apicomplexa also arose multiple times independently.

To further investigate the convergent evolution of parasitic lifestyles in *Piridium* and *Platyproteum*, we examined plastid retention and function, a well-studied trait of the Apicomplexa (*2, 3*). With both genomic and transcriptomic data from *Piridium*, we first assembled its complete plastid genome (Fig. 2A), which is strikingly similar in size, architecture, and gene content to apicoplast genomes (Fig. 2B). The *Piridium* plastid genome is a highly reduced compact circle (∼34kb) with all remaining genes in perfectly synteny with homologues in its closest relative, the photosynthetic *Vitrella.* Similar to the apicoplast, it is extremely AT-rich (21% G+C content), and uses a non-canonical genetic code where UGA encodes tryptophan (as seen in *Chromera, Toxoplasma* and Corallicola, but not in the more closely related *Vitrella*) (*14, 15*). It retains similar ribosomal genes as well as the same bacterial RNA polymerases (rpoB, rpoC1, rpoC2) and other protein-coding genes (sufA, clpC, tufA) as apicoplasts. It has also convergently lost all genes relating to photosynthesis, as well as rps18, rpl13, rpl27, secA and secY (Fig 2C). Reflecting its origin from a chrompodellid ancestor, the *Piridium* plastid also encodes a handful of genes that are present in *Vitrella* but absent from apicoplasts: rps14, rpl3, and rpoA. Curiously, only a partial rRNA inverted repeat remains in *Piridium*; this repeat is ancestral to all apicomplexans and chrompodellids, but has also similarly been lost in the piroplasm apicomplexans, *Babesia* and *Theileria*(*16, 17*).

**Fig. 2:**
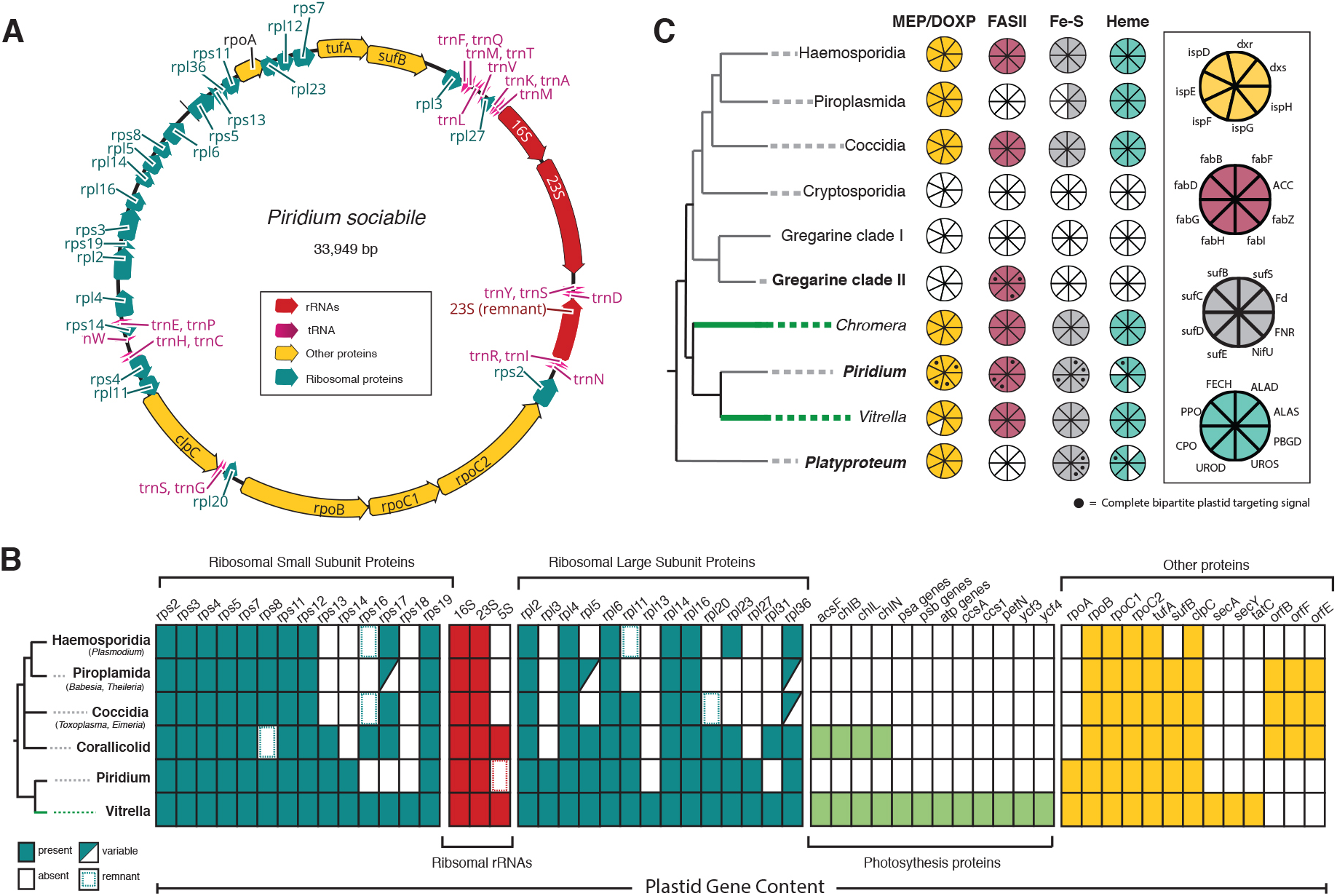
Plastid dependency in *Piridium* and *Platyproteum* has evolved convergently to apicomplexans. (A) Complete annotated plastid genome of *P. sociabile.* (B) Presence of plastid biosynthetic pathways across the tree of apicomplexans and chrompodellids. Portions of the circles represent the proteins found in each pathway found (key shown on right). Black circles indicate the presence of complete N-terminal bipartite plastid targeting peptides (only shown for newly added transcriptomes) (C) Plastid gene content of apicomplexans and *Vitrella* (free-living, photosynthetic) compared to *Piridium*.

Apicomplexans depend on apicoplasts for essential biosynthesis of four compounds: isoprenoids (the MEP/DOXP pathway), heme, iron-sulfur (Fe-S) clusters, and fatty acids (the FASII pathway) (*2*). All apicomplexans rely on these pathways except piroplasms, which have lost the FASII pathway and use cytosolic FASI instead, and *Cryptosporidium*, which can salvage the metabolites from its host and has lost its plastid entirely (*18, 19*). We identified all enzymes from these pathways, and all enzymes for analogous and homologous cytosolic pathways using profile hidden Markov models (HMMs) and analyzed the resulting genes for evidence of distinctive N- terminal bipartite plastid targeting peptides (Fig. 2B). It is impossible to conclude that any single gene is absent based on transcriptomic data alone, so only the absence of all genes for entire biochemical pathways is considered here. The dependency on plastid metabolism in *Piridium* is identical to most apicomplexans, with the retention of all four pathways but no photosystems or other known plastid functions. *Platyproteum* is similar, but has also lost the FASII pathway, and so more resembles the piroplasms *(17,18)*.

Interestingly, the same analysis on the clade of gregarines revealed a greater degree of variation from other apicomplexans than seen in the cryptic plastids that evolved in parallel. Like *Cryptosporidium*, the terrestrial gregarines *Monocystis* and *Gregarina* have completely lost all plastid metabolism and likely also lost the organelle (which also suggests that the phylogenetic relationship between *Cryptosporidium* terrestrial gregarines remains uncertain)(*19,20*). In contrast, however, the marine gregarines *Lecudina* and *Pterospora* retain the complete FASII pathway, but no other identifiable plastid metabolism. This is the first evidence of a plastid in any gregarine and is also functionally curious since it is isoprenoid biosynthesis that has been proposed to be the main barrier to plastid loss(*3*). The gregarines suggest that plastid dependency is highly context-dependent.

Looking beyond the plastid, metabolic reconstructions based on KEGG identifiers across the whole genome confirm an overall convergence of functional reduction, but also some divergence (Fig. 3). Both *Piridium* and *Platyproteum* have, as expected, substantially reduced their metabolic functions compared with free-living chrompodellids. However, neither is as reduced as apicomplexan parasites. In both cases a few core pathways such as the glyoxylate cycle and pyrimidine catabolism have been retained. Of the two, *Piridium* contains the greatest breadth of biosynthetic functions that were mostly lost in all other parasitic groups, such as *de novo* amino acid biosynthesis (isoleucine and arginine) and purine biosynthesis (inosine) and degradation. Surprisingly, however, the gregarine *Monocystis agilis* has also retained a greater metabolic capacity than other apicomplexans, revising the baseline metabolic complexity of the group as a whole.

**Fig. 3:**
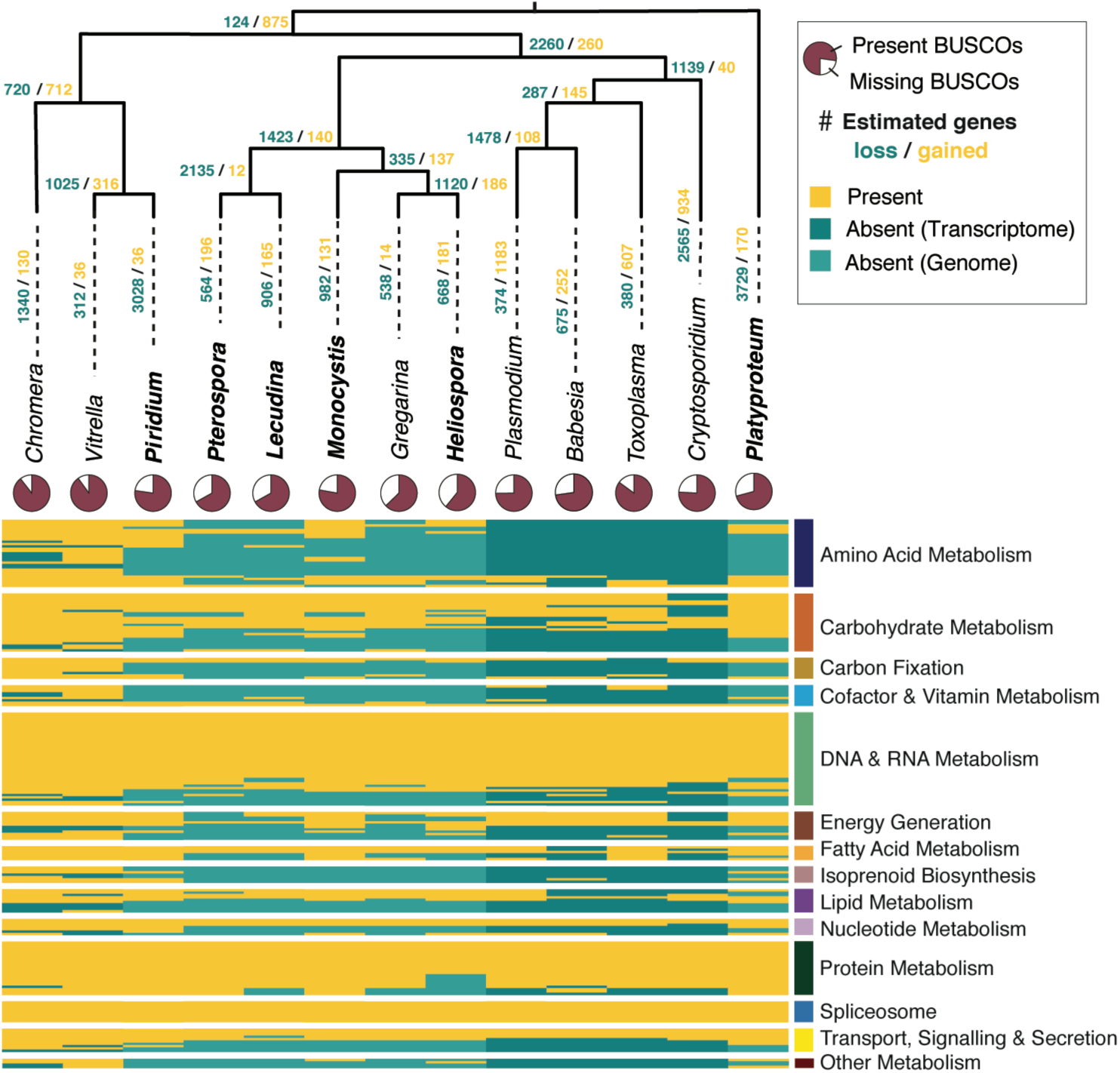
The distribution of cellular metabolic pathways across the tree of apicomplexans and chrompodellids. The list of metabolic pathways are shown on the right. Yellow represents presence and shades of blue indicate absence based on genomic data (dark blue), or absence based on transcriptomic data (lighter blue). Our newly sequenced transcriptomes are shown in bold lettering. Estimated gains and losses of genes (orthogroups) are shown on nodes and on the branches leading to each species. The pie charts show the percentage of genome or transcriptome completeness based on BUSCO scores.

The origin of apicomplexan parasites from free-living photosynthetic alga represents a major evolutionary transition between two very different modes of living, so different in this case that the idea was originally met with considerable skepticism. The current data show that, however dramatic this transition may seem, it was not unique, but rather repeated at least three times in related lineages of photosynthetic algae. The convergent forms of parasitism in *Piridium, Platyproteum*, and apicomplexans, suggest that the ancestors of these lineages maintained high levels of redundancy in metabolic pathways between compartments that persisted over long periods of evolutionary time, and apparently shared some predilection to animal parasitism. The underlying reason for this is not clear, since the evolution of apicomplexan parasitism is not linked to the acquisition of any novel feature or machinery, but is instead marked by loss and tinkering of the existing genomic repertoire.

## Acknowledgments

We thank Maria Herranz, Niels van Steenkiste, Phil Angel and the Hakai Institute for their assistance in sample collection. Also, Filip Husnik for his guidance with the transcriptomics, and Waldan Kwong for his valuable feedback on the manuscript.

## Authors contributions

V.M., M.K. and P.J.K. designed the study. **V**.M., ÁK., M.A.F., B.S.L. obtained samples. M.A.F. and Á.K. performed the initial molecular diagnosis for *Piridium*. V.M., M.K. and N.A.T.I. conducted genomics, transcriptomics and phylogenomics. L.H., N.A.T.I., M.K. and V.M. performed plastid analyses. V.M. and M.K. performed all other analyses. V.M., M.K. and P.J.K. wrote the manuscript with input from all authors.

## Data and materials availability

The transcriptome and genome survey reads have been deposited in the NCBI Sequence Read Archive (PRJNA539986). The *Piridium* plastid genome has been deposited in GenBank (accession no. SUB5567202). Single gene trees used in the phylogenomic and plastid analyses were deposited in DataDRYAD repository.

