## Supplementary Materials for "Multiple independent origins of apicomplexan-like parasites"

**This PDF file includes:**

Materials and Methods

Figs. S1 to S2

Tables S3 to S5

Materials and Methods

*Piridium* isolation and DNA/RNA extraction:

Common whelks, *Buccinum undatum*, were collected using dredges across Breidafjördur, West coast of Iceland, 65° 7.576'N; 22° 44.738'W. Whelks were sedated using 0.1% MgSO_4_ in seawater for 1-2 hours, then examined for the presence of *Piridium* cysts on the surface of the foot using a dissection microscope. Mature (large) cysts were gently squeezed with pointed forceps until the *Piridium* gamonts were released. The resulting exudate was collected into concave glass spot plates containing filtered seawater and rinsed with autoclaved seawater three times to remove host tissues and mucous. DNA and RNA from the resulting gamonts was then extracted using a QIAGEN, Allprep DNA/RNA Mini Kit (Cat. No. 80204).

*Piridium* transcriptome and genome library preparation, sequencing & transcriptome assembly:

Piridium cDNA was synthesized using the SMARTseq2 protocol with seven cDNA amplification cycles(*21*). RNA and DNA sequencing libraries were both prepared using Illumina Nextera XT and Nextera protocol respectively, and sequenced using 2 × 300bp Illumina MiSeq (DNA) and 2x100bp Illumina HiSeq 2000 run (RNA). Both RNA and DNA reads were adapter and quality trimmed with Trimommatic (*22*). RNA reads were further processed to remove low complexity regions using PRINTseq (*23*) and were assembled into transcripts using Trinity v2.4 (with default settings) and translated into protein sequences using Transdecoder v5 *(24,25)*.

Assembly and annotation of *Piridium* plastid genome:

The MIRA4 assembler was used to assemble the genomic DNA reads, which led to the assembly of single circular plastid genome chromosome (*26*). This assembly was validated by mapping of the reads back to the assembly by Bowtie2 (*27*). The plastid genome was then automatically annotated using MFAnnot (http://megasun.bch.umontreal.ca/cgi-bin/mfannot/mfannotInterface.pl) and RNAweasel (http://megasun.bch.umontreal.ca/cgi-bin/RNAweasel/RNAweaselInterface.pl), followed by manual corrections in Geneious v11.1.5 (https://www.geneious.com).

Host collection, single-cell isolation and transcriptomics of gregarines and *Platyproteum*:

*Monocystis agilis* was isolated from the seminiferous vesicles of earthworms (*Lumbricus terrestris*), purchased from Berry's Bait and Tackle, Richmond, British Columbia, Canada. *Lecudina tuzetae* and *Heliospora caprellae* were isolated from the guts of the animals *Platyneries bicanaliculata,* and *Caprella californica* respectively. They were both collected at low tide from Calvert Island, BC, Canada. *Pterospora schizosoma* was isolated from the gut of a bamboo worm, *Axiothella rubrocincta,* that was collected from Friday Harbour, Washington, USA. *Platyproteum* sp. was isolated from the gut of a Sipunculid worm that was collected from Sint Joris Bay, Curaçao.

All parasites were collected in the trophozoite life stage (large feeding cells). Trophozoites were released into filtered seawater by teasing apart the intestines/seminal vesicles of the respective hosts with pointed forceps. The trophozoites were washed at least three times in autoclaved filtered seawater, or ultrapure water (for *Monocystis*) and viewed and photographed under a Leica DMIL LED microscope equipped with a 40× objective and a Sony α6000 camera. Single trophozoite cells were picked using a glass capillary micropipettes and transferred to a 0.2 mL thin-walled PCR tube containing 2 µL of cell lysis buffer (0.2% Triton X-100 and RNase inhibitor (Invitrogen)). cDNA was synthetized from the single cell, or a pool of 2-3 cells, using the Smart-Seq2 protocol(*1*). The cDNA concentration was quantified on a Qubit 2.0 Fluorometer (Thermo Fisher Scientific Inc.).

Prior to high-throughput sequencing, 1µL of the final cDNA product was used as a template for a PCR amplification of the V4 region of the 18S rRNA gene using Phusion High-Fidelity DNA Polymerase (New England Biolabs, Thermo Scientific) and the general eukaryotic primer pair TAReuk454FWD1 and TAReukREV3 (*28*). The PCR product was then sequenced by Sanger dideoxy sequencing. The SSU rRNA gene sequences were used to confirm the identity of the newly collected organisms and avoid animal host contamination using BLASTn to look for similar sequences in the non-redundant NCBI database (*29*). Once the identity of the parasite was confirmed, sequencing libraries were prepared using the Nextera XT protocol, and sequenced on a single lane of Illumina MiSeq using 250 bp paired end reads. Raw reads are available in the NCBI Short Read Archive (SRA) (PRJNA539986).

Transcriptome assembly of gregarines and *Platyproteum:*

The raw Illumina sequencing reads were merged using PEAR v0.9.6, and FastQC was used to assess the quality of the paired reads (*30*, *31*). The adapter and primer sequences were trimmed using Trimmomatic v0.36 and the transcriptomes were assembled with Trinity v2.4.0 (*22*, *24*). The contigs were then filtered for animal host contaminants using BlobTools in addition to blastn and blastx searches against the NCBI nt database and the Swiss-Prot database, respectively (*29*, *32*). Coding sequences were predicted using a combination of TransDecoder v3.0.1 and similarity searches against the Swiss-Prot database (*25*). Assessment of the quality of the assembly and annotation of the transcriptomes (including *Piridium*) was carried out using BUSCO (*33*).

Ortholog identification, gene concatenation and phylogenomics:

In addition to our newly generated transcriptomes, the following transcriptomes and genomes were downloaded from EuPathDB and screened for orthologs; Hammondia hammondi, Sarcocystis neurona, Eimeria falciformis, and Gregarina niphandrodes (34). All transcriptomes were comprehensively searched for a set of 263 genes that have been used in previous phylogenomic analyses (*12,13*). All the sequences in the 263 gene-set, representing a wide range of eukaryotes, were used as queries to search the above datasets using BLASTn (*29*). The hits were then filtered using an e-value threshold of 1e-20 and a query coverage of 50%. Each of the gene-sets was then aligned using MAFFT L-INS-i v7.222 and trimmed using trimAl v1.2 with a gap-threshold of 80% (*35, 36*). Single gene trees were then constructed to identify paralogs and contaminants using IQ-TREE v.1.6.9 (LG+G4 model) or RAxML v8.2.12 (PROTGAMMALG model) with support from 1000 bootstraps (*37,38*). The resulting trees were manually scanned in FigTree v1.4.2 and contaminants and paralogous sequences were identified and removed (*39*). The final cleaned gene-sets were filtered so that they contained only a maximum of 40% missing OTUs and then concatenated in SCaFoS v1.2.5 (*40*). The resulting concatenated alignment consisted of 198 genes spanning 58,116 amino acid positions from 39 taxa. The phylogenomic maximum likelihood tree was constructed with the heterogenous mixture C40+LG+Γ4+F model as implemented in IQ-TREE (model LG+Γ4+F yielded identical topology) (*38*). Statistical support was inferred using 500 bootstrap replicates using the LG+C40+Γ4+F PMSF profiles, and 1000 bootstrap replicates using the LG+Γ4+F model in RAxML (*37,41*). The Bayesian tree was computed using Phylobayes (*42*) under the GTR-CAT model with constant sites removed from the analyses. Four independent chain were run for 9 thousand generations and converged with maxdiff = 0.19 (20% burning) (Fig. 1G).

For the plastid based phylogenomic analyses, a previously published dataset *(43)* was used and enriched with proteins from wide sets of publicly available apicoplast proteins and the *Piridium* plastid genome. The plastid phylogenomic tree was constructed using a concatenated alignment of 62 plastid-encoded proteins with RAxML using the LG+Γ4+F substitution model with 500 bootstrap replicates (Fig. S2).

Search for plastid proteins:

Profile hidden Markov models (HMMs) were used to identify plastid metabolic proteins in our transcriptomes based on curated alignments. To construct the curated alignments, known dinoflagellate proteins were used as queries in a BLASTp search (e-value threshold of 1e-5) against a comprehensive custom database containing representatives comprised of major eukaryotic groups, with a focus on plastid-containing lineages (dinoflagellates, chrompodellids, Apicomplexa, cryptophytes, haptophytes, stramenopiles, Archaeplastida) as well as selected taxa from non-plastid lineages (Opisthokonta, Amoebozoa, Apusozoa, Ancyromonadida and ciliates) and RefSeq data from all bacterial phyla at NCBI (https://www.ncbi.nlm.nih.gov/,

last accessed December 2017) *(19*). The database was subjected to CD-HIT with a similarity threshold of 85% to reduce redundant sequences and paralogues *(44)*. Results from blast searches were parsed for hits with a minimum query coverage of 50% and e-values of less than 1e-25 (or 1e-5 for HemD). The number of bacterial hits was restrained to 20 hits per phylum (for FCB group, most classes of Proteobacteria, PVC group, Spirochaetes, Actinobacteria, Cyanobacteria (unranked) and Firmicutes) or 10 per phylum (remaining bacterial phyla) as defined by NCBI taxonomy. Parsed hits were aligned with MAFFT v. 7.212, using the --auto option, poorly aligned regions were eliminated using trimAl v.1.2 with a gap threshold of 80% *(35,36*). Maximum likelihood tree reconstructions were then performed with FastTree v. 2.1.7 using the default options *(45)*. The resulting phylogenies and underlying alignments were inspected manually to remove contaminations, recent paralogs and duplicate sequences. The cleaned, unaligned sequences were then subjected to filtering with PREQUAL using the default options to remove non‑homologous residues introduced by poor-quality sequences, followed by alignment with MAFFT G‑INS‑i using the VSM option (‑‑unalignlevel 0.6) to control over-alignment *(36,46*). The alignments were subjected to Divvier (https://github.com/simonwhelan/Divvier) using the ‑divvygap option to improve homology inference before removing ambiguously aligned sites with trimAl v. 1.2 (gap threshold of 1%)*(35*). Trees for final sequence curation were calculated with IQ-TREE v. 1.6.5, using the ‑mset option to restrict model selection (to DAYHOFF, DCMUT, JTT, WAG, VT, BLOSUM62, LG, PMB, JTTDCMUT) for ModelFinder, while branch support was assessed with 1000 ultrafast bootstrap replicates, and once more subjected to manual inspection *(38,47)*.

Profile HMMs were then generated using these curated alignments and HMM searches were conducted on all transcriptomes and genomes using HMMER v3.1 and an e-value threshold of 1e-5 *(48)*. All the hits were then extracted and incorporated into the original alignments and realigned using MAFFT v7.222 (--auto option). The resulting alignments were then used to generate phylogenies in IQ-TREE v.1.6.9 using the LG+F+G4 substitution model and statistical support was assessed using 1000 ultrafast bootstrap replicates (*38,47*). The phylogenies were then manually scanned in FigTree v1.4.2 and contaminants, paralogs, mitochondrial sequences, and long-branching divergent sequences were identified and removed(*39*). The remaining sequences were realigned and used to generate maximum likelihood phylogenies in IQ-TREE v.1.6.9 (*38*). Phylogenetic models were selected for each tree individually based on Bayesian Information Criteria using ModelFinder as implemented in IQ-TREE, and statistical support was assessed using 1000 ultrafast bootstrap pseudoreplicates (*29*, *49*).

Search for plastid localization signals:

To investigate the N-terminal extensions and thus intracellular location of proteins of interest, corresponding alignments were manually inspected for completeness of the sequences and for N‑terminal extensions relative to prokaryotic or cytosolic homologs. Prediction of signal peptides as part of N-terminal bipartite leader sequences was performed with the Hidden Markov Model of SignalP3.0 using the default truncation setting of 70 residues (*50*). To predict putative N‑terminal transmembrane domains, TMHMM v. 2.0 was used only on the first 100 amino acid residues of the transcript to improve prediction accuracy (*51*). Putative plastid transit peptides were interpreted as 24-aa stretches downstream of the signal peptide, representing the minimum length for apicomplexan transit peptides still within the N-terminal extension and upstream of the estimated start of the mature protein, as described by Parsons et al. *(52).* Conserved domains and their coordinates in the mature protein region of candidate sequences were identified with the Pfam sequence search service on http://pfam.xfam.org/search/sequence, using the gathering threshold as a cut-off (Table S1).

Mapping cellular metabolic pathways:
We reconstructed the metabolic maps for our new transcriptomes, as well as representative species across the apicomplexan and chrompodellids, using Kyoto Encyclopedia of Genes and Genomes (KEGG) *(53)*. We first assigned KEGG ortholog identifiers (KO) to all proteomes using the web-based server, KAAS (KEGG Automatic Annotation Server) (http://www.genome.jp/kegg/kaas/), and where possible we used annotations already available within KEGG. The assigned KO numbers were used to identify complete metabolic pathways using the KEGG reconstruct module and module mapper. Complete metabolic pathways present in *Piridium, Platyproteum* or both but missing in other apicomplexans were further investigated. The identity of all proteins in these unique pathways were confirmed using BLASTp to ensure removal of contaminants and false positives (Table S2).

Ortholog identification and search for apicomplexan invasion/extracellular proteins:

Orthofinder was used to infer orthologs within the apicomplexans, chrompodellids and *Platyproteum*, while *Oxytricha*, *Tetrahymena*, *Symbiodinium* and *Perkinsus* were used as a outgroup for the analyses *(54)*. Diamond searches were used to identify homologues between pairs of taxa *(55)*. The Orthofinder results were then analyzed using Dollo parsimony as implemented in the program Count to obtain estimates of gene gain and loss across different species *(56)*. Previously published sets of invasion (98 proteins from *Plasmodium falciparum* 3D7) and extracellular (722 proteins from diverse apicomplexans) proteins were then identified within orthogroups and their orthologs in the studied taxa were recorded (Table S3) *(4)*.

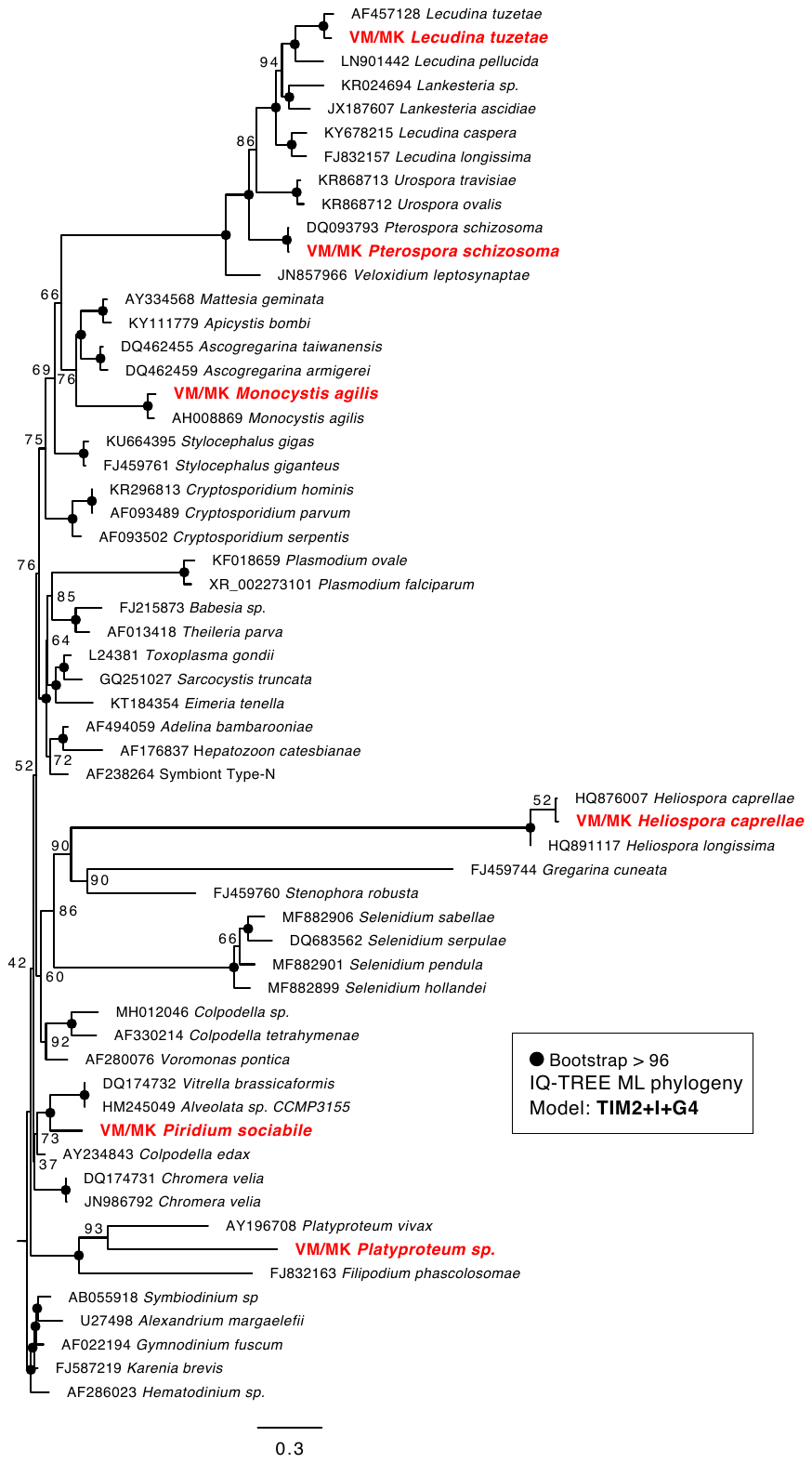

**Fig. S1.** Myzozoan 18S SSU rRNA gene phylogeny inferred from Maximum Likelihood (TIM2+I+G4 IQ-TREE substitution model) with 1000 ultrafast bootstraps (UFB) for support. The tree includes a diversity of apicomplexans, chrompodellids and a dinoflagellate outgroup. Taxa denoted in red represent 18S sequences retrieved from our sequenced transcriptomes. Black dots on nodes denote UFB > 96 and all other support values are written on the node.

­­

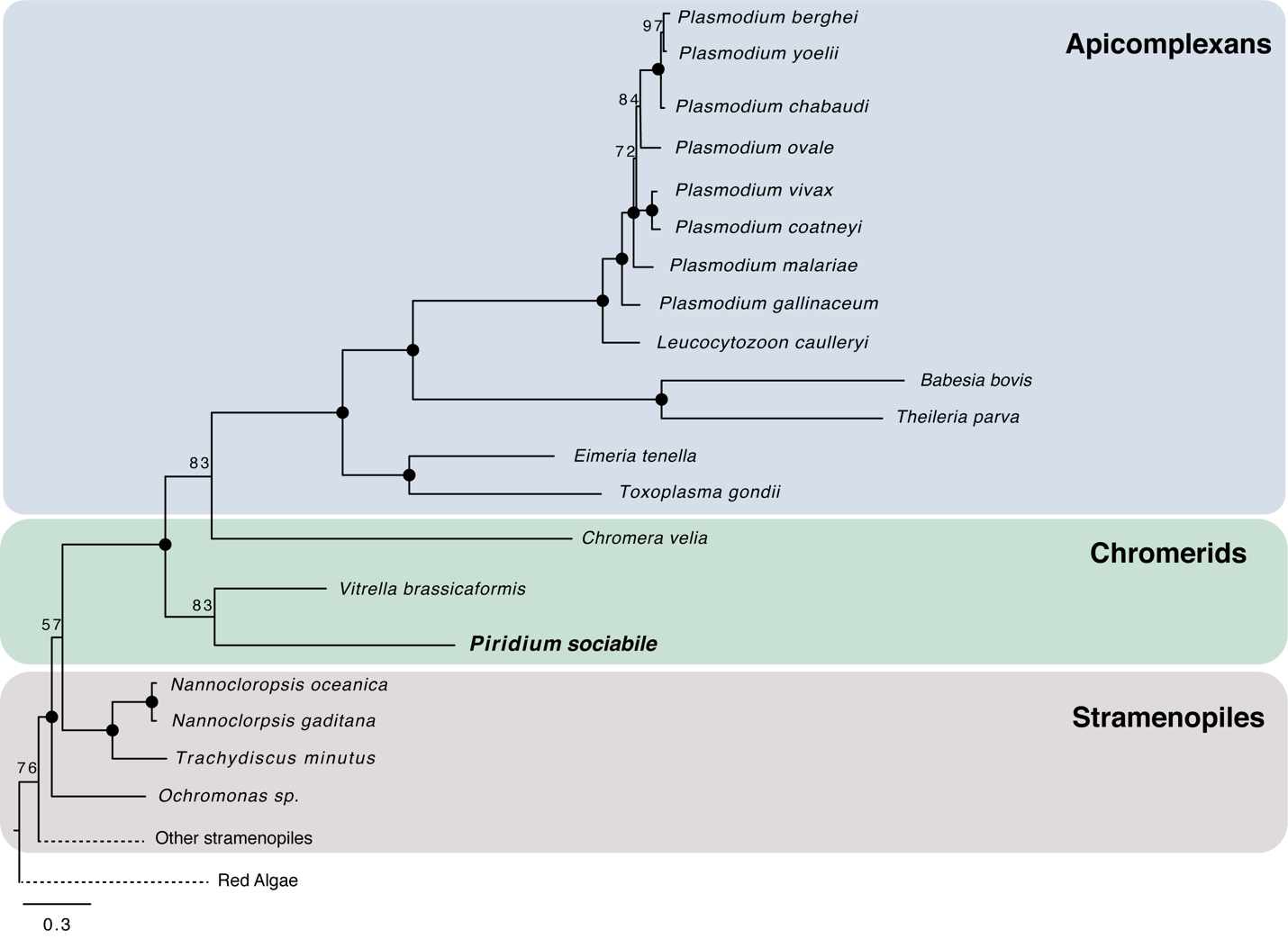

**Fig. S2:** Maximum likelihood tree based on 62 plastid genome encoded proteins, constructed in RAxML with the LG+GAMMA+F model of sequence evolution. Statistical support was inferred from 500 bootstrap replicates (BP) using the same model. Black dots denote 100% BP support.

Table S1.

Plastid-Targeted Proteins and Localization Signal Analysis.

**Table S2.**

Presence of Metabolic Pathways based on KEGG Modules.

Table S3.

Orthologs of proteins involved in apicomplexan infection (invasion and extracellular proteins).
